# SCAMPR: Single-Cell Automated Multiplex Pipeline for RNA Quantification and Spatial Mapping

**DOI:** 10.1101/2022.03.23.485552

**Authors:** Ramin Ali Marandi Ghoddousi, Valerie M. Magalong, Anna K. Kamitakahara, Pat Levitt

## Abstract

Spatial gene expression, achieved classically through *in situ* hybridization, is a fundamental tool for topographic phenotyping of cell types in the nervous system. Newly developed techniques allow for the visualization of multiple mRNAs at single-cell resolution, greatly expanding the ability to link gene expression to tissue topography. Yet, methods for efficient and accurate quantification and analysis of high dimensional *in situ* hybridization are limited. To this end, the Single-Cell Automated Multiplex Pipeline for RNA (SCAMPR) was developed, facilitating rapid and accurate segmentation of neuronal cell bodies using a dual immunohistochemistry-RNAscope protocol and quantification of low and high abundance mRNA signals using open-source image processing and automated segmentation tools. Proof of principle using SCAMPR focused on spatial mapping of gene expression by peripheral (vagal nodose) and central (visual cortex) neurons. The analytical effectiveness of SCAMPR is demonstrated by identifying the impact of early life stress on differential gene expression by vagal neuron subtypes.

**Motivation:** Quantitative analysis of spatial mRNA expression in neurons can lack accuracy and be both computationally and time intensive. Existing methods that rely on nuclear labeling (DAPI) to distinguish adjoining cells lack the precision to detect mRNA expression in the cytoplasm. In addition, quantification methods that rely on puncta counts can generate large, variable datasets that potentially undercount highly expressed mRNAs. To overcome these methodological barriers, we developed the SCAMPR pipeline that allows for fast, accurate segmentation of neuronal cell body boundaries, topographic gene expression mapping, and high dimensional quantification and analysis of mRNA expression in tissue sections.

## Introduction

The murine brain contains approximately 70 million individual neurons (Herculano-Houzel et al., 2006). Understanding a tissue of such complexity requires knowledge about its component parts, both at the level of single neurons and at the level of cell groups that constitute structural and functional nodes in circuits. Analysis of the individual characteristics of single neurons, such as their topographical organization, morphology, and molecular signature (among other properties), can be used to categorize them into distinct functional cell groups (Erö et al., 2018; Zeng & Sanes, 2017). The commercialization and ongoing development of single molecule fluorescent *in situ* hybridization (smFISH) techniques (e.g. RNAscope, HCR RNA-FISH, MERFISH, etc.), supplemented by a variety of image analysis tools and software that quantitate molecular signal with cellular specificity, allow for accessible multimodal comparison of transcriptomics and histology at a single-cell level (Chen et al., 2015; Choi et al., 2018; Eng et al., 2019; Goh et al., 2020; Ståhl et al., 2016; F. Wang et al., 2012). Existing methods for quantitative analysis of smFISH face several challenges, including incomplete single-cell segmentation and a loss of cell size information due to the use of the nuclear marker DAPI to delineate cell boundaries, limited compatibility with multi-round versions of smFISH that are required for analysis of more than 4 genes, and resource/time intensive pipelines (Carine Stapel et al., 2016; Codeluppi et al., 2018; Dries et al., 2019; Maynard et al., 2020; Y. Wang et al., 2021). Here, we introduce SCAMPR, a quantitative analysis pipeline that combines commercialized smFISH (HiPlex RNAscope) with immunohistochemistry (IHC), and employs existing and newly developed semi-automated image processing tools for accurate segmentation and quantification of mRNA expression in neurons (F. Wang et al., 2012). SCAMPR includes open-source code for quantitative analysis and spatial mapping of mRNA expression in neurons, as well as precise comparison of gene expression between distinct experimental groups. In addition, neuron size is preserved accurately due to whole-soma segmentation, and SCAMPR generates datasets that require relatively small amounts of digital storage and processing power for downstream analysis. Lastly, the entirety of the SCAMPR pipeline, from the benchwork to the generation of quantitative plots, takes 5 days to complete when multiplexing 12 genes, making it possible to perform comparative, multimodal gene expression experiments in a short amount of time.

We validate the compatibility of SCAMPR on two mouse nervous system structures that have distinct neuronal packing densities and topographical organizations: the nodose ganglion (NG) of the peripheral nervous system and the primary visual cortex (V1) of the central nervous system. In V1, we demonstrate that SCAMPR can be used to correlate single and multidimensional gene expression to cortical layer topography and to distinguish cell types based on soma size, gene expression, and location. We use SCAMPR in the NG to demonstrate cell-type specific gene expression changes resulting from early life stress (ELS). Our findings demonstrate the accuracy and utility of SCAMPR for descriptive and comparative analysis of neuronal gene expression and topography.

## Results

### Overview of SCAMPR

A comprehensive snapshot of the SCAMPR pipeline is provided, outlining the steps involved in processing and analyzing images from a single tissue section (Figure 1). SCAMPR has two workflows: 1) an assay workflow (Figure 1A), which comprises benchtop *in situ* hybridization, immunostaining, microscopy, and image pre-processing, and 2) an image processing and data analysis workflow (Figure 1B), which includes cell segmentation, additional batch image processing in FIJI/ImageJ, mRNA quantification, and data analysis (Schindelin et al., 2012; Schneider et al., 2012). The SCAMPR assay workflow (Figure 1A) starts with the published protocol for the Advanced Cell Diagnostics (ACD) HiPlex RNAscope Assay (F. Wang et al., 2012). Briefly, probes targeting three distinct mRNAs are hybridized to the tissue, each with different fluorescent signal, a DAPI counterstain is performed, the tissue is imaged, the fluorophores are cleaved, and this process is repeated with the hybridization of three new mRNA probes. This is repeated in the same tissue until all genes of interest have been hybridized and imaged. If infrared confocal imaging is available, four probes can be hybridized simultaneously. After all probe signals have been imaged, DAPI counterstaining and HuC/D immunochemistry are performed to mark the boundaries of cellular nuclei and the neuronal cytoplasm, respectively, and the tissue is imaged one final time. Flattened, maximum intensity projection images from the different hybridization/staining rounds are spatially registered (overlayed) using the DAPI signal with the ACD RNAscope HiPlex Image Registration Software or in ImageJ. The resulting registered images then proceed through the SCAMPR Image Processing & Data Analysis Workflow (Figure 1B).

**Figure 1.**
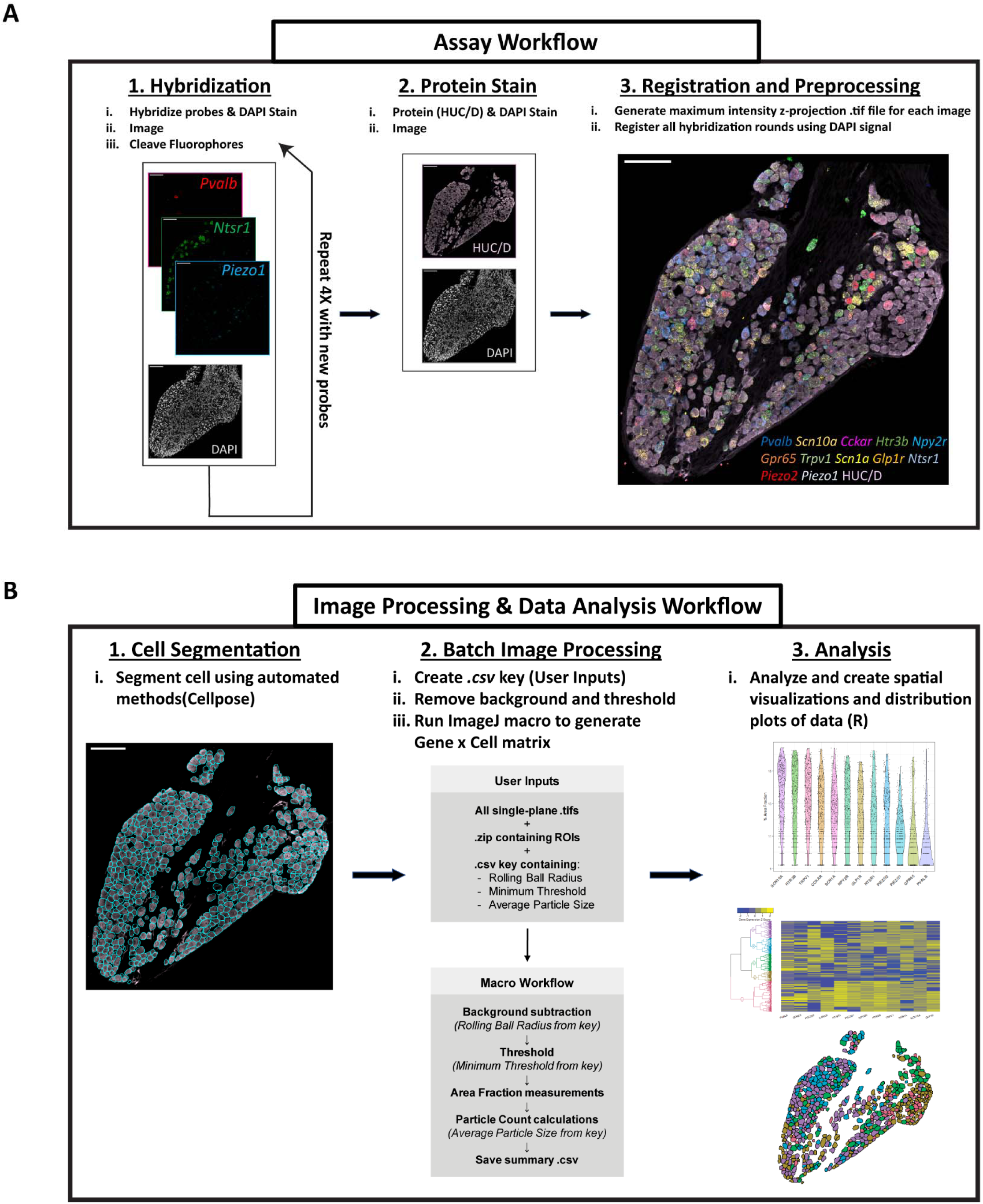
Schematic of SCAMPR Pipeline. **A.** Assay workflow: 1. All rounds of HiPlex RNAscope are performed and sections are imaged. 2. HuC/D immunohistochemistry is performed subsequently. 3. Images from all rounds are pre- processed and registered using the DAPI signal as the anchor. **B.** Image processing and data analysis workflow: 1. Batch cell boundary segmentation using Cellpose. 2. Images are inspected by the user then processed using macros to automate gene expression quantification for each cell in ImageJ. 3. Gene expression (top) and co-expression, clustering (middle), and spatial mapping (bottom) are performed in R. Scale bars = 100 um.

In this second workflow (Figure 1B), cell segmentation and batch image processing are performed. The HuC/D image is used to segment the cells into individual regions of interest (ROIs). These ROIs may be achieved by manual drawing in FIJI/ImageJ, or to save time, by using an open-source automated cell segmentation program such as Cellpose (Stringer et al., 2021). In parallel, the images containing the mRNA signal are visually inspected by the user in ImageJ to determine the optimal parameters for subtracting background and for applying a threshold to convert the mRNA signal into a binary signal/no-signal image. The ROIs are then used in combination with these parameters to quantify signal for each mRNA probe on a single cell level in each tissue section. New macros were written to automate this process in ImageJ. These macros produce a “Gene by Count” matrix in the form of a CSV file, which is used for subsequent data analyses. The data analysis component of the SCAMPR pipeline contains numerous methods to quantify gene expression, co-expression, and global expression patterns, as well as methods for spatial mapping and comparative gene expression analysis between groups. Each analysis method has its own R script that can be accessed on GitHub.

### Dual IHC-RNAscope accurately demarcates cell boundaries in peripheral and central nervous system tissues

HuC and HuD are mRNA-binding proteins that are involved in transcript regulation, and are selectively expressed in neurons starting in embryonic development, initially appearing in progenitor cells, and continuing to be expressed in postmitotic neurons into adulthood (Okano & Darnell, 1997). HuC and HuD are present at the protein level in both the nucleus and cytosol (Diaz-Garcia et al., 2021). The whole-cell expression, neuron specificity, and broad expression time course of HuC/D make them excellent cell boundary markers for quantitative studies that require neuronal segmentation. Two experiments employing SCAMPR demonstrate that HuC/D immunohistochemistry is compatible in multiple nervous system tissues at different developmental timepoints. First, multi-round HiPlex RNAscope is successfully applied in postnatal day 9 (P9) nodose ganglion (NG) of the peripheral nervous system as well as in adult primary visual cortex (V1) in the central nervous system (Figures 2 A, B, E, and F). Subsequent HuC/D immunostaining is compatible with HiPlex RNAscope in these same tissues and accurately labels neuronal cell bodies following multi-round HiPlex RNAscope (Figures 2 C and G). Comparison of HuC/D labeling to nuclear DAPI labeling demonstrates that HuC/D more precisely captures the complete individual cellular profile than the nuclear DAPI signal, resulting in accurate localization of single-cell RNAscope signals in both tissues (Figures 2 C & D, and G & H). Additional validation of RNAscope-HuC/D co-staining was performed in the embryonic NG and early postnatal brainstem (data not shown). In all instances, intense HuC/D signal allowed for distinct segmentation between individual neurons. There is minimal immunolabeling of neuronal processes, further enhancing the ability to distinguish between closely apposed neuronal somata.

**Figure 2:**
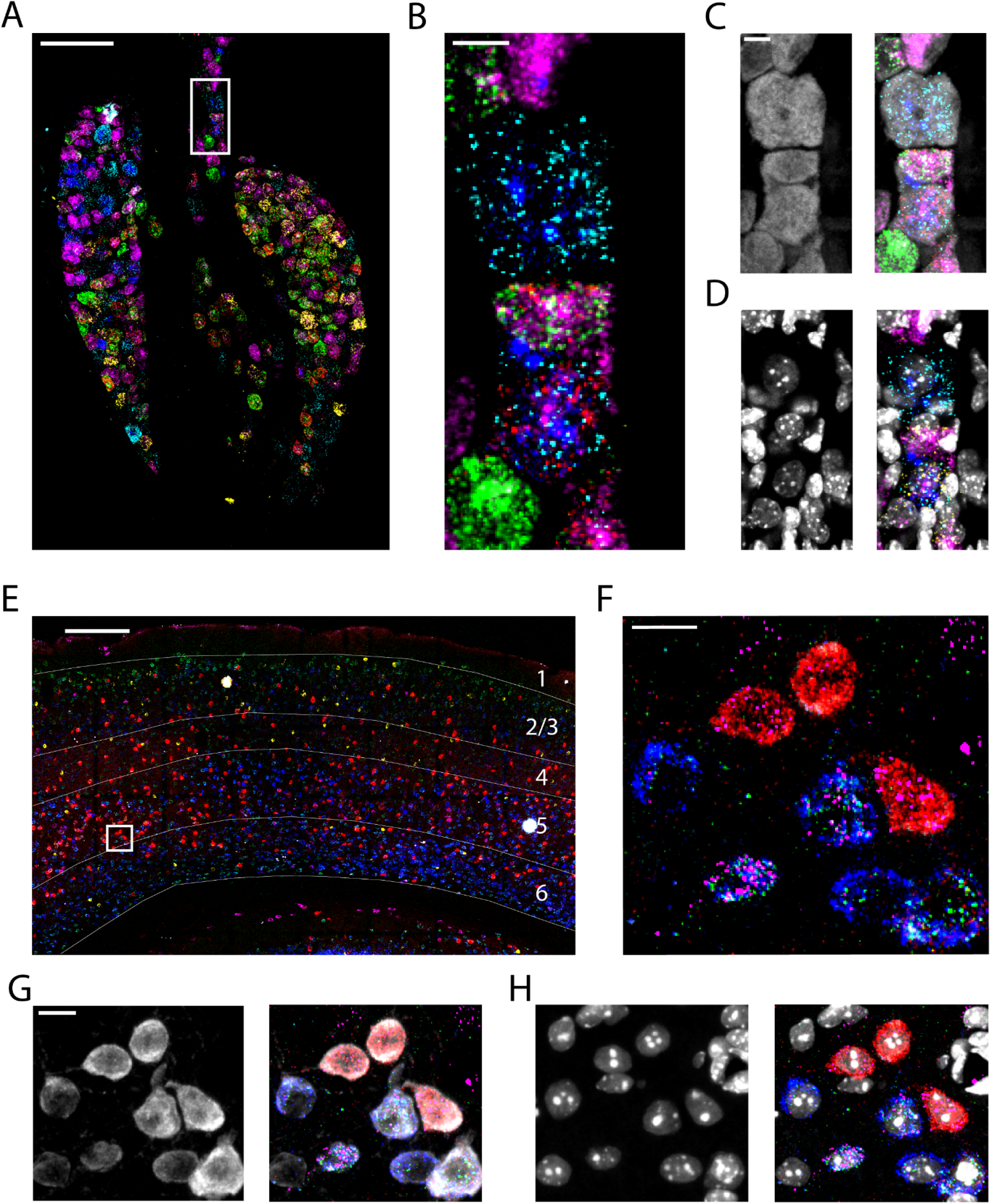
Combined IHC and HiPlex RNAscope allows for high resolution, single cell mRNA expression in the central and peripheral nervous system. **A.** HiPlex RNAscope visualized in postnatal day 9 nodose ganglion. Scale bar = 100 um. **B.** High magnification inset from A. Scale bar = 15 um. **C and D.** High magnification insets from (A) co-visualized with the cytosolic marker HuC/D (C) and nuclear marker DAPI (D). Scale bar = 10 um. **E.** HiPlex RNAscope visualized in adult mouse visual cortex. Scale bar = 100 um **F.** High magnification inset from (E). Scale bar = 5 um. **G and H.** High magnification inset from (E) co-visualized with cytosolic marker HuC/D (G) and nuclear marker DAPI (H). Scale bar = 5 um.

### Automated methods for accurate segmentation of single neurons

Quantitative analysis of spatial transcriptomics data requires segmentation of cells into single, discrete units. The most accurate segmentation of cells in a tissue section will be obtained with more precise cell boundary markers that demarcate the cytoplasm, rather than using a single organelle label such as the nucleus. DAPI marks cell nuclei, and because mRNA is localized to both nuclear and cytosolic cell compartments, segmentation based on the nucleus can result in mRNA fluorescent signal that is unassigned to a cell profile and therefore not measured. Using expanded nuclear labeling (cytoDAPI) allows for more accurate segmentation of whole neurons but makes it difficult to separate neurons from non-neurons in densely packed tissues with large amounts of non-neuronal cell types (Y. Wang et al., 2021). Using the localization of two genes, manual segmentation was performed to demonstrate potential differences in mRNA signal detection between nuclear segmentation using DAPI signal and whole-cell segmentation using HuC/D staining after HiPlex RNAscope in a sample section of the V1 cortex (Figure 3A). The HuC/D signal allows for segmentation of the cell body, and more accurately encircles the entirety of the mRNA signals (here *Pvalb* and *Chd13)* in the segmented cells compared to the DAPI signal (Figure 3B).

**Figure 3:**
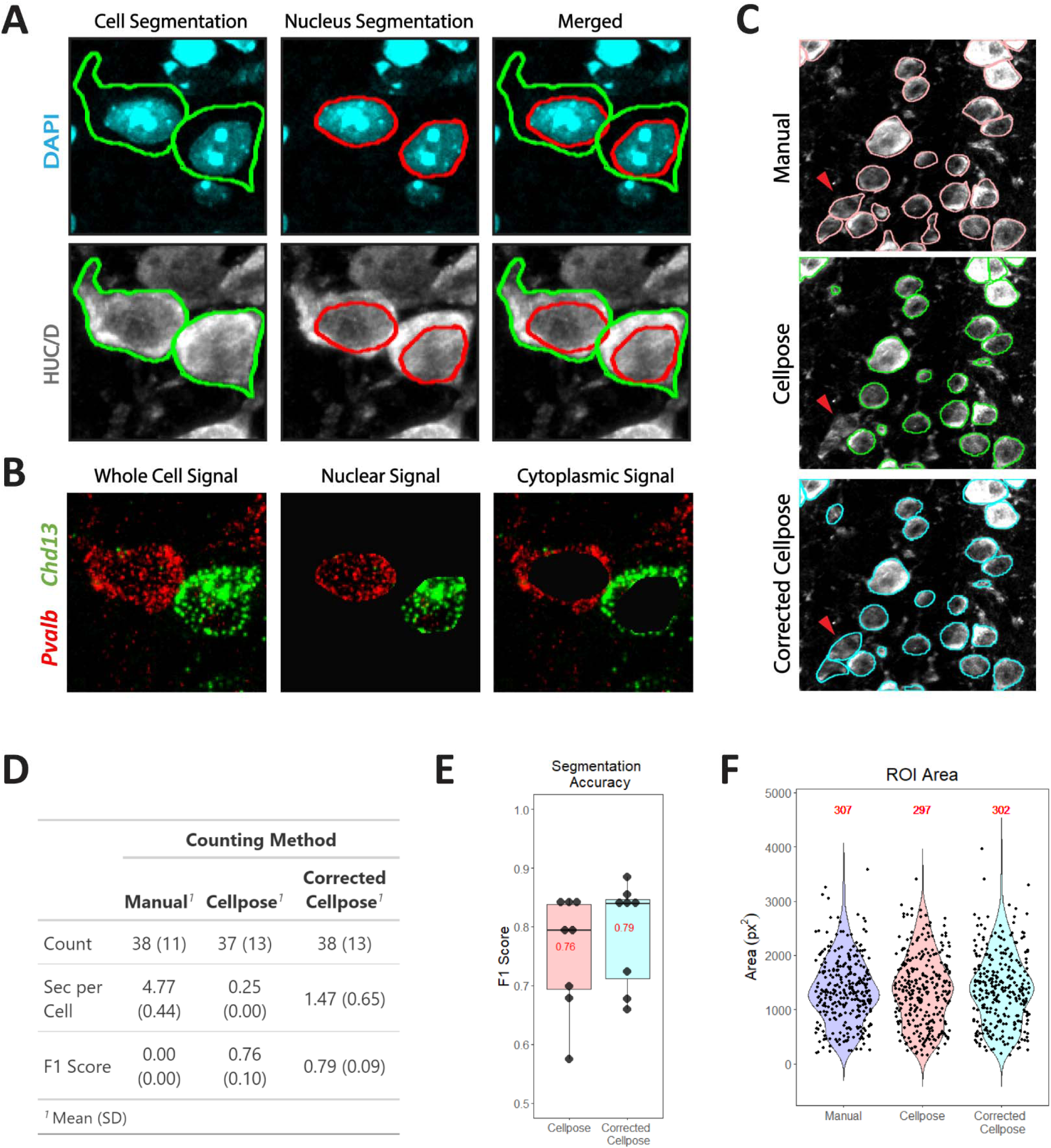
Automated segmentation of cytoplasm of single cells using Cellpose. **A.** Differences in nuclear segmentation boundaries using DAPI (red outline) and whole-soma segmentation boundaries using HuC/D (green outline) in V1 cortex. These segmented boundaries serve as regions of interest (ROI) for quantitative analyses. **B.** Nuclear signal that is present in the cytoplasm for two genes is not included when using nuclear segmentation. **C.** Visualization of whole-cell segmentation using the HuC/D signal. Automated Cellpose segmentation is compared to manual segmentation and a mix of Cellpose and manual segmentation (Corrected Cellpose). Red arrowhead denotes a cell that was identified during manual segmentation but not during automated Cellpose segmentation. **D.** Mean cell counts, time burden, and F1 scores for manual, Cellpose, and Corrected Cellpose segmentation methods. n = 8 images per group. **E.** Mean and individual F1 scores for ROIs generated by Cellpose and Corrected Cellpose segmentation as compared to the ground truth ROIs generated by manual segmentation. n = 8 images per group. **F.** The total number of ROIs generated after Cellpose or Corrected Cellpose segmentation were within 3% of the total number of ROIs generated Manual segmentation (see total number of cells in red). Cell area distributions are nearly indistinguishable between the three methods, demonstrating high fidelity and accuracy of automated whole-cell Cellpose segmentation on HuC/D expressing neurons. n = 8 images per group.

The time requirement for manual segmentation increases dramatically as the surface area of the tissue of interest and the cell packing density increases. Popular automated segmentation methods such as watershed segmentation greatly reduce this time requirement but perform poorly on images of densely packed cell bodies, a particular challenge in most developing and certain adult tissues. More recent methods such as StarDist are trained on specific image sets (DAPI signal) and require further user-guided training to accurately define whole-cell boundaries that are more closely apposed (Weigert et al., 2020). Cellpose, a generalized, deep learning, automated segmentation algorithm that has been trained on multiple different image sets, obviates these challenges (Stringer et al., 2021). Application of Cellpose on HuC/D immunolabeling signal produces highly comparable results to manual segmentation in the same tissue sections. Small deviations (missed cells, incorrect boundaries, etc.) can be corrected manually after Cellpose segmentation (Corrected Cellpose) (Figure 3C). On a per cell basis, data from two different operators demonstrates that there is an approximately 20-fold difference in segmentation time between using Cellpose and manual segmentation, with Cellpose segmentation requiring 0.25 seconds per cell and manual segmentation requiring 4.77 seconds per cell. Corrected Cellpose takes ∼6 times longer than Cellpose alone but is also ∼3 times faster than manual segmentation (Figure 3D). The calculation of an F1 score, representing the accuracy of Cellpose and Corrected Cellpose segmentation as compared to manual segmentation, demonstrates the precision of this segmentation method (See Methods for details). To generate the F1 score, ROIs that showed a ≥70% overlap with the manually generated ROIs were considered true positives (Caicedo et al., 2019). This comparison resulted in an F1 score of 0.76 for Cellpose and 0.79 for Corrected Cellpose (Figures 3D and E). To better understand whether this accuracy affected the overall dynamics of segmentation in each image, the number of segmented cells and the distribution of ROI areas were compared between each method. Cellpose and Corrected Cellpose resulted in only 1.6-3.0% fewer ROIs than manual segmentation, and the distribution of the ROI areas of the neurons between the three methods were nearly identical, providing further evidence for the overall accuracy of the fully automated segmentation method compared to the manual method (Figure 3F). These data demonstrate that Cellpose segmentation alone can be employed for both high segmentation accuracy and substantial time-savings.

### Quantification and spatial projection of pairwise mRNA expression patterns

One of the main advantages of SCAMPR is the ability to **accurately** quantify multiplexed fluorescent *in situ* signal on a single cell level while preserving the topographic location of each cell in the tissue. The pipeline provides an opportunity to analyze both average gene expression and expression topography with single cell spatial resolution across all neurons in a single tissue section. Single neuron expression of nine different genes was quantified in V1 (Figure 4A). The analysis of V1 not only reveals differences in mean gene expression between transcripts (e.g. *Slc17a7* vs *Chat*), but also demonstrates that for some genes with similar mean expression (*Chd13* vs *Piezo1*), the neurons have distinct expression distribution ranges. Specifically, *Piezo1* expression is normally distributed and exhibits a smaller distribution range while *Chd13* expression is skewed leftward towards zero and exhibits a larger overall range of expression (Figure 4A).

**Figure 4:**
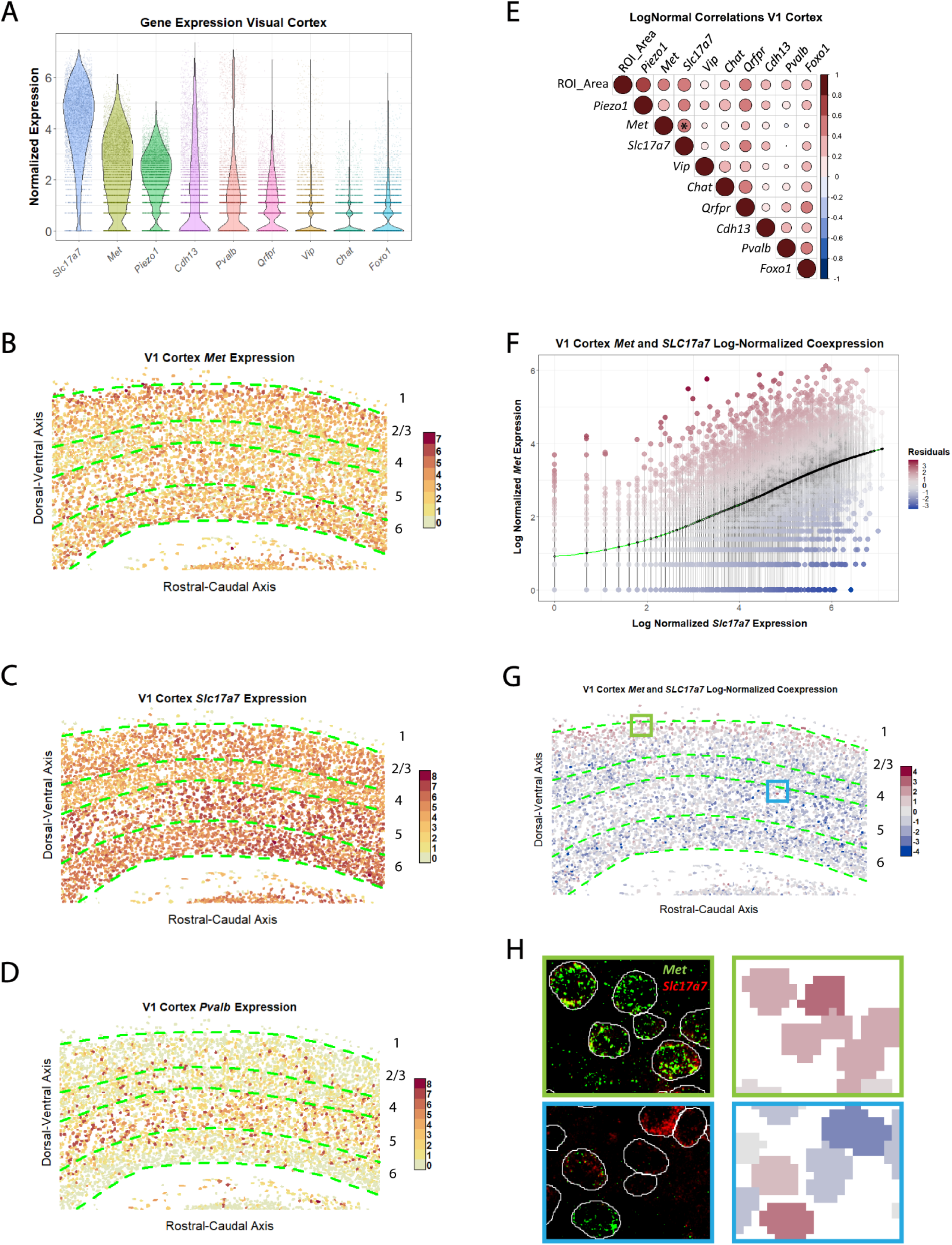
Analysis and spatial visualization of gene expression and co-expression of HiPlex RNAscope. **A.** Violin plot of log-normalized gene expression for nine genes expressed in V1 cortex. **B.** Spatial map of log-normalized *Met* expression in the V1 cortex. Note the layer specific differences in expression (greater expression in layers 2-3 shown in red, and low expression in layer 4 shown in yellow). **C.** Spatial map of log-normalized *Slc17a7* (encoding the vesicular glutamate transporter) expression in V1 cortex. Note the greater expression in layers 5 and 6 and medium-low expression in layer 4. **D.** Spatial map of log-normalized *Pvalb* expression in V1 cortex. Note high-expressing red profiles scattered across all layers, with an enrichment in layer 5. **E.** Pairwise correlation plot for log-normalized gene expression in V1 cortex. Asterisk denotes correlation between *Met* and *Slc17a7* which is analyzed further in F and G. **F.** LOESS regression plot between log-normalized *Met* and *Slc17a7* expression in V1 cortex. Points denote single cells and are colored based on their distance from the regression line (residual size). **G.** Spatial map of residuals from F. Red cells have large positive residuals and express more *Met* and/or less *Slc17a7* than is predicted by the model. Blue cells have large negative residuals and express less *Met* and/or more *Slc17a7* than is predicted by the model. Note layer 2 enrichment of red cells. **H.** Left: High magnification micrographs, denoted by green (layer 2) and blue (layer 4-5 border) boxes in G. Top image illustrates the relatively high expression of *Met* (green) compared to *Slc17a7* (red) in layer 2. The resulting large positive residuals (pink/red) are shown in the top right graphic. Bottom image illustrates cells with a high *Slc17a7* (red) in layer 4, lower co-expression in one profile and higher expression of *Met* (green) in a layer 5 neuron. These expression patterns match with their residual-denoted colors in the spatial map (bottom right).

SCAMPR also facilitates the mapping of gene expression and gene co-expression patterns at single-cell spatial resolution onto the tissue of origin. This is demonstrated by using SCAMPR to map 3 genes, *Met*, *Slc17a7*, and *Pvalb,* each having distinct levels of expression and neuronal distributions in V1 cortical layers. Consistent with prior results from our laboratory, neurons expressing the highest levels of *Met* are distributed predominantly in layer 2 and those expressing the lowest levels of *Met* localize to layer 4 (Figure 4B) (Eagleson et al., 2016; Judson et al., 2009). *Slc17a7* is expressed in neurons across all layers in V1, with slightly lower expression in layer 4 (Figure 4C).

Neurons expressing *Pvalb* at high levels also are scattered across all layers of V1, with noticeable enrichment in layer 5 (Figure 4D). In addition to single gene spatial mapping, SCAMPR also can be used to quantify and spatially map gene co-expression patterns. Gene co-expression was quantitated by generating pairwise linear correlation coefficients between all nine genes in V1. A variety of relations are revealed, with some genes highly co-expressed (*Slc17a7* and *Piezo1*), and others discordant (*Slc17a7* and *Pvalb*) (Figure 4E). As an added dimension, the accurate segmentation of neuron cell bodies, rather than nuclei, provides cell size (ROI_Area) as a variable in this co-expression analysis. Matching prior reports in AMPA and NMDA gene expression in motor neurons, the quantitative expression levels of most genes in V1 exhibited positive correlations with cell area (Rana et al., 2020). One obvious exception was noted, with *Vip* exhibiting expression levels that were independent of cell size (Figure 4E).

A unique feature of data analysis using SCAMPR is the single-cell, topographic characterization of gene co-expression patterns between two genes. This is accomplished by fitting a trend line to the co-expression scatterplot of the two genes, and subsequently measuring residual distances between each cell and the trend line. Here, the residual distances were mapped back onto the cells in the tissue. Fitting a locally estimated scatterplot smoothing (LOESS) line to the *Met* and *Slc17a7* co-expression scatterplot shows a positive relationship between *Met* expression and *Slc17a7* expression for the large majority of *Met/Slc17a7* co-expressing cells. Cells that fall on or near the trend line represent the majority or “expected” co-expression pattern for *Met* and *Slc17a7*, whereas those that fall far from the trend line express relatively more or relatively less *Met* and *Slc17a7* than expected. On the tissue section rendition, these subtypes were marked in increasingly darker shades of red if they expressed more *Met* and/or less *Slc17a7* than expected, and increasingly darker shades of blue if they expressed less *Met* and/or less *Slc17a7* than expected (Figure 4F). The positive (red), negative (blue), or neutral (grey) distances of these cells from the trend line were then mapped back onto their location in the V1 tissue, showing that the *Met*/*Slc17a7* ratio was larger than the trend in layer 2 and smaller than the trend in multiple layers, particularly in layer 6a (Figure 4G). To validate the accuracy of this method for characterizing both trends and thus, the heterogeneity in the co-expression pattern of two genes, high-magnification insets from layers 2 and 5 of the spatial map were compared to the original HiPlex RNAscope images from the same regions (Figure 4H).

### Clustering and spatial mapping of high dimensional mRNA expression patterns

In addition to single and dual gene expression patterns, SCAMPR can utilize hierarchical clustering methods to categorize neurons into groups based on global gene expression patterns. Using nine genes assayed in the V1 cortex HiPlex RNAscope experiments, SCAMPR was used to cluster the cells into eight distinct groups. The clusters are represented as a joint dendrogram/heatmap (Figure 5A). Because ROI identity and spatial coordinates are preserved during the clustering, each cluster can be spatially mapped back onto the tissue section. Notably, some of the clusters segregate spatially. Cells that comprise cluster 1 are predominantly located in layer 4, whereas cells in cluster 8 are predominantly located in layers 5 and 6a (Figure 5B). Additionally, the cells that comprise cluster 8 are larger than those in cluster 1 (Figure 5C). Cells in cluster 8 also exhibit either higher or equivalent log normalized gene expression when compared to cells in cluster 1, with prominent differences in genes such as *Cdh13* and *Foxo1* (Figure 5D). Examination of the original images for mRNA fluorescence validates significantly higher *Cdh13* and *Foxo1* expression in deeper compared to superficial layers, with the highest co-expression in layer 5 and a near absence of these two genes in layers 2-4 (Figure 5E).

**Figure 5:**
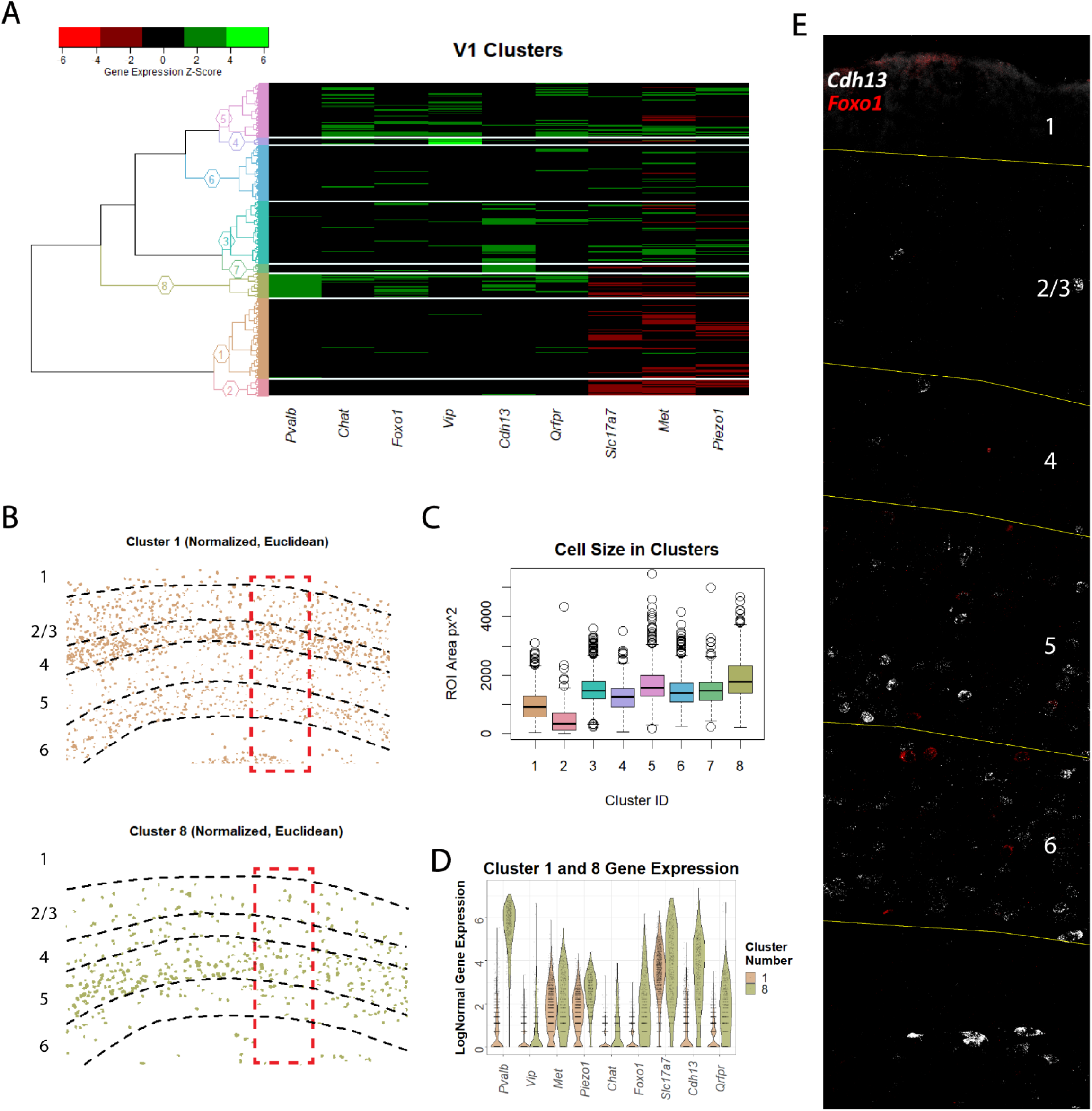
High-dimensional clustering analysis of HiPlex RNAscope data. **A.** Heatmap of scaled gene expression in V1 cortex after hierarchical clustering. Cluster numbers shown on y-axis. **B.** Spatial map of cells in Cluster 1, which are mostly localized in layers 2/4/6, and Cluster 8, which are predominantly localized in layers 5/6a. High resolution images of red boxes presented in E. **C.** Boxplots demonstrating differences in cell size (area) between the clusters. Largest difference in cell size is between Cluster 1 (brown) and Cluster 8 (green). **D.** Violin plots demonstrating higher expression of several genes in Cluster 1 compared to Cluster 8. **E.** Hiplex RNAscope image of *Chd13* (white) and *Foxo1* (red), which are enriched in Cluster 8 and absent in Cluster 1. Note that both genes are more highly expressed in layers 5 and 6a, the primary location of Cluster 8 cells.

### Using SCAMPR to identify gene expression pattern differences between two experimental groups

Differences in expression levels of specific genes due to a manipulation may either vary across all neurons that express a specific transcript, or in specific subtypes, and can be resolved using quantitative analyses at the single cell level. Thus, in addition to its utility in spatial genomics, high dimensional *in situ* hybridization can be used to compare the expression of a carefully selected set of genes, such as cell subtype markers, to determine potential phenotypic differences that may occur due to an experimental manipulation between groups. Here, we used a well-validated model of limited bedding and nesting (LBN), applied from postnatal day (P)2-9 in mice. LBN disrupts maternal care and produces an early-life-stress (ELS) response in pup offspring (Eagleson et al., 2020; Heun-Johnson & Levitt, 2016; Rice et al., 2008). Care-as-usual (CAU) pup offspring were raised under normal conditions. Development of vagal circuitry is sensitive to ELS, though information is limited at a molecular level (Banihashemi & Rinaman, 2010; Card et al., 2005). To investigate prospective ELS-induced vagal gene expression changes in early postnatal mice, we used SCAMPR to analyze the expression of a set of 12 genes that demarcate adult subtypes of vagal sensory neurons located in the nodose ganglion (NG) of ELS and CAU mice (Kupari et al, 2019). Tissue was harvested for processing on postnatal day (P) 9 at the end of the 7-day period of ELS exposure. Pairwise correlation coefficients and least squares regression analyses were performed for both groups separately to identify possible gene co-expression differences between groups (Supplemental Figure 1). Visual inspection of the plot matrix was sufficient to narrow possible differences in gene co-expression between CAU and ELS. Analyses began with the identification of relatively well-correlated gene pairs for all cells from all groups (*R* > 0.40). From these, gene pairs that had **<u>both</u>** prominent between-group differences in correlation coefficients and linear regression line slopes were selected. Two gene pairs fit these criteria: *Scn10a/Ntsr1* and *Scn1a/Pvalb* (Supplemental Figure 1). To identify differences between the co-expression of these genes in CAU and ELS groups, scatterplots of *Scn10a/Ntsr1* co-expression and *Scn1a/Pvalb* co-expression were generated, and LOESS models were fitted for each group. These plots demonstrated lower *Pvalb* and *Ntsr1* expression with rising values of *Scn10a* and *Scn1a* in the ELS group when compared to the CAU group (Figure 6A and 6B). Since *Scn10a* and *Scn1a* have been shown to be broad cell-type markers in the NG, these analyses at single-cell resolution suggest that there are subtype-specific decreases in *Pvalb* and *Ntsr1* expression in NG cells after early life stress exposure (Kupari et al., 2019).

**Figure 6:**
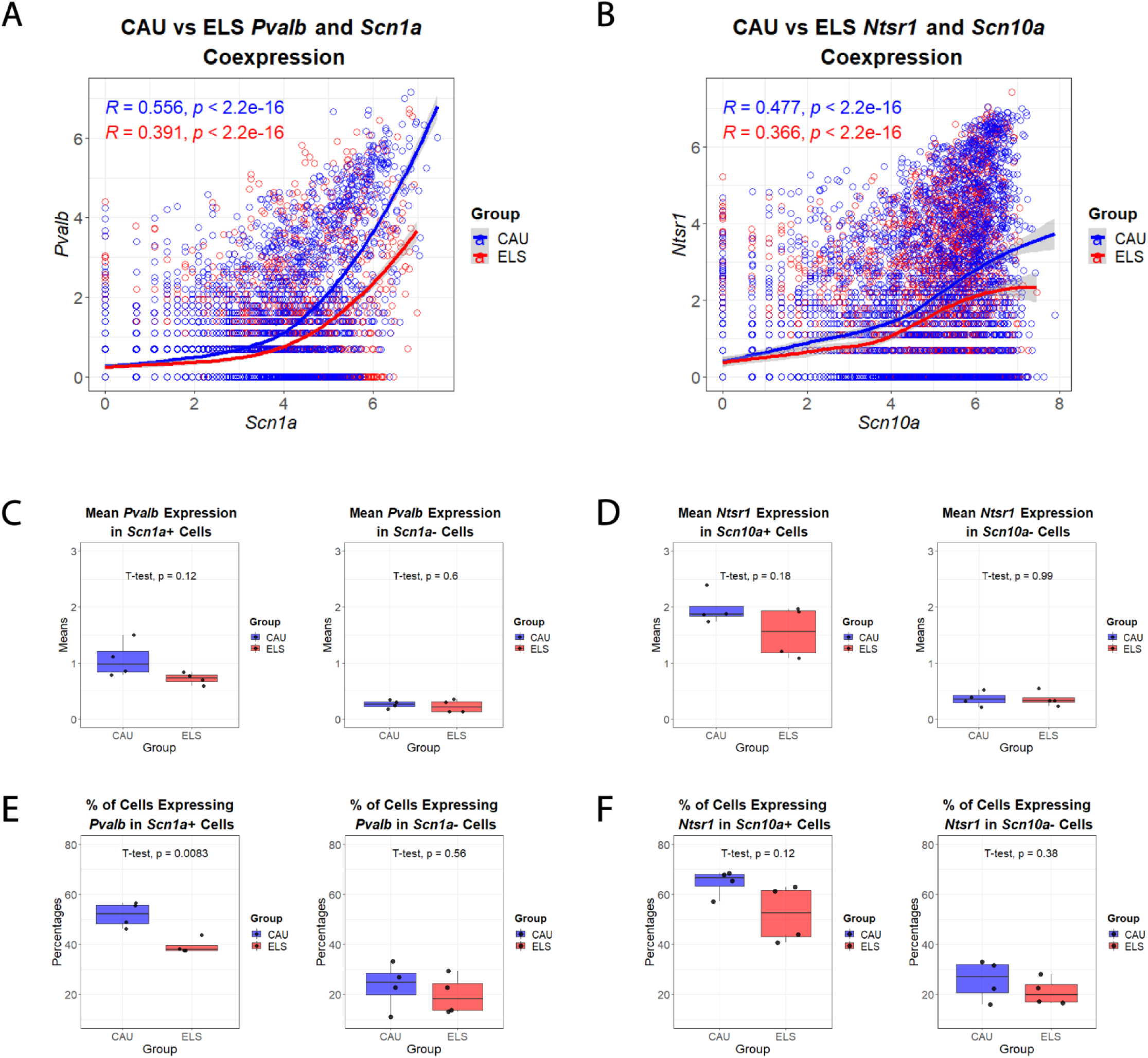
Utilization of HiPlex RNAscope for comparison of gene expression at single-cell resolution in nodose ganglion neurons from CAU and ELS experimental groups. **A and B.** *Pvalb*/*Scn1a* and *Ntsr1*/*Scn10a* co-expression scatterplots for all cells in the dataset. Points denote single cells in CAU (blue) and ELS (red) mice. Solid blue and red lines denote the LOESS lines of CAU and ELS mice, respectively. Pearson’s r and corresponding p-values for the correlations were determined for each group separately. **C.** Mean, per-animal *Pvalb* expression in ELS and CAU *Scn1a*-expressing (left) and non *Scn1a*-expressing (right) cells. (n = 4 per group). No statistically significant difference noted between groups. **D.** Mean, per-animal Ntsr1 expression in ELS and CAU *Scn10a*-expressing (left) and non *Scn10a*-expressing (right) cells. (n = 4 per group). No statistically significant difference noted between groups. **E.** Percentage of cells from CAU and ELS animals that express *Pvalb* in *Scn1a*-expressing (left) and non *Scn1a*-expressing cells (right). (n = 4 per group). Note the statistically significant decrease in the percentage of neurons expressing *Pvalb* in *Scn1a*-positive neurons in the ELS mice. **F.** Percentage of cells from CAU and ELS animals that express *Ntsr1* in *Scn10a*-expressing (left) and non *Scn10a*-expressing cells (right). (n = 4 per group). No statistically significant difference noted between groups.

The subtype-specific reductions observed could be driven either by an average decrease in expression of *Pvalb or Ntsr1* across many cells, or a decrease in the proportion of cells that express each gene. To investigate these possibilities, *Scn1a-*expressing (*Scn1a*+), *Scn10a*-expressing (*Scn10a*+), non-*Scn1a*-expressing (*Scn1a*-), and non-*Scn10a*-expressing (*Scn10a*-) data subsets were generated and mean *Pvalb* and *Ntsr1* expression was calculated for NG neurons from each ELS and CAU pup. The averages of these mean expression values were compared between groups for each subset. There was no difference in mean *Pvalb* and *Ntsr1* expression in the *Scn1a*- and *Scn10a*-subsets of neurons, and while mean *Pvalb* and *Ntsr1* expression in ELS mice trended towards a decrease in in the *Scn1a*+ and *Scn10a*+ subsets of neurons, these differences did not reach statistical significance (Figure 6C and D).

It has been shown that the proportion of cells expressing a particular gene can be altered in an experience-dependent manner during development (Cheng et al., 2022). To determine whether the reduction of *Pvalb* and *Ntsr1* expression in ELS animals reflects a reduction in the proportion of cells expressing the genes rather than overall mean expression levels in the specific subpopulations, the percent of cells expressing *Pvalb* and *Ntsr1* in the different cell-type specific subsets was quantitated (Figure 6E and 6F). In *Scn1a*+ cells, there was a significant 24.1% decrease in the percentage of cells expressing *Pvalb* in ELS animals when compared to CAU animals (51.8% CAU vs 39.3% ELS, p ≤0.05), whereas in *Scn1a-* cells, there was no significant difference between groups (23.5% CAU vs 19.7% ELS, p = 0.56) (Figure 6E). In both *Scn10a*+ and *Scn10a*-cells, there was a trending decrease in *Ntsr1*-expressing cells in ELS animals when compared to CAU animals (64.7% CAU vs 52.2% ELS, p = 0.12 and 25.8% CAU vs 21.1% ELS, p = 0.38) (Figure 6F). These analyses suggest that in ELS animals, *Pvalb* expression is reduced in a subset of NG cells that express *Scn1a* and that this reduction is partially driven by a reduction in the fraction of cells expressing *Pvalb* transcript. In addition, broad gene expression comparisons of the remaining ten genes demonstrated a decrease in mean *Gpr65* expression in ELS animals, yet no difference in the percentage of cells expressing this gene (Supplemental Figure 2). Together, these experiments demonstrate that SCAMPR can be used as a tool to quantitatively compare global and cell-type specific differences at single cell spatial resolution, as well as gene co-expression patterns across experimental groups.

## Discussion

Here we present SCAMPR, a novel, open-source, user-friendly, time-saving, bench-to-desk pipeline for accurate quantification and analysis of spatial neuronal gene expression with single-cell resolution. The quantification and analytical tools that comprise SCAMPR allow for comparison of the expression patterns of multiple genes, spatial mapping of relative gene expression and co-expression patterns across a tissue, and cell-type specific comparisons of gene expression between experimental groups, all at single neuron resolution. The comparative analyses of cell labeling methods demonstrated superior savings of time, fully completing bench and analytical work in approximately 5 days, without sacrificing accuracy, while using fully automated or semi-automated segmentation of the entire neuronal soma.

### Novel methods enabling accurate spatial analyses at single neuron resolution

Utilization of HuC/D staining in the SCAMPR pipeline allows for demarcation of neurons across multiple developmental timepoints and multiple subregions of the nervous system that exhibit different cell packing densities and spatial organization patterns, allowing broad application of SCAMPR for analyzing topographic gene expression in neurons. The stability of the anti-HuC/D antibody under the harsh protease digestions and antigen retrieval processing steps of HiPlex RNAscope allows for HuC/D staining at the very end of hybridization protocol, circumventing the early occupation of an imaging channel or the need for protein stain degradation/quenching or fluorophore cleaving. Other sufficiently stable primary antibodies marking astrocytes, microglia, oligodendrocytes that are present in the nervous system, or other antibodies marking non-nervous system cell classes that are present in other tissue types can be validated for tissue processing stability and used with SCAMPR to quantify gene expression differences in myriad cell types. SCAMPR utilizes a combination of Cellpose and ImageJ scripts to quantify gene expression for each neuron. In contrast to methods that perform puncta counting, which rely on intense particles standing out from their surroundings, SCAMPR employs a detection-by-thresholding method to quantify the number of pixels in a binary image that are positive for a fluorescent signal representing a particular gene. Because adjoining fluorescent molecules do not need to be separated from one another based on intensity in detection-by-thresholding (contrasting to puncta counting methods, where such molecules might be unreliably quantified), this method allows for accurate quantification of genes expressed at low or high levels within the same tissue section or even in adjoining neurons.

Furthermore, SCAMPR quantifies gene expression using flattened, z-projected images, and outputs gene expression as the percentage of the cell surface area or the number of pixels within a cell area that are positive for a gene. This contrasts with some puncta count methods which match puncta with cellular ROIs by assigning x, y, and z coordinates to each individual puncta, thus generating large data matrices that increase the computational burden in the requisite downstream analyses.

This is an important feature when considering data management and computational time savings, given that the hybridization and imaging technology facilitates coverage of larger tissues containing more cells. Lastly, segmentation based on the use of both DAPI and HuC/D, in conjunction with modified versions of the ImageJ quantification macros, can be used to obtain cell compartment information by separately quantifying nuclear and cytosolic mRNA signals.

### Spatial mapping tools enable visualization of complex expression, co-expression, and cell-type clustering patterns

The capability of SCAMPR for fast and accurate segmentation of neurons and quantification of *in situ* hybridization signal were combined with its tools for spatial analysis of gene expression, co-expression, and clustering driven cell-type localization in the V1, a highly topographic structure with a laminar organization. We posited that challenging the ability of SCAMPR to accurately segment, measure, and map gene expression in a highly organized brain structure that is characterized by region-specific differences in packing density would reveal strengths and weaknesses of the pipeline. The data demonstrate the capabilities of SCAMPR for detecting layer-enriched differences in the expression of single genes, two genes, and multiple genes in this brain region. Novel analyses of the co-expression of two genes (*Met* and *Slc17a7*), in which each cell was binned based on residual proximity to a fitted line, facilitated spatial mapping of the general co-expression patterns of these genes in the majority of neurons. Using this same analysis method, SCAMPR was able to extract minority populations of neurons with more unique co-expression patterns based on their deviation from the general co-expression patterns in the majority population of neurons. This holistic analysis method will compliment more direct strategies of separating neuron populations based on gene co-expression patterns, such as the semi-quantitative categorization of neurons based on the expression levels of both genes (low-low, low-high, high-low, and high-high), or by considering their expression distributions and summary statistics (min, max, mean, quartiles, etc). Lastly, it is possible to integrate HiPlex RNAscope and other smFISH data with published single cell RNA sequencing datasets (Hashikawa et al., 2020; Y. Wang et al., 2021). These integrated datasets could theoretically be used with SCAMPR to spatially map the expression levels of genes from single cell RNA sequencing experiments.

### SCAMPR enables analysis of differentially expressed genes in subsets of cell types

SCAMPRs utility for comparing gene expression differences between neurons developing under two different environmental conditions was demonstrated. In the NG, a tissue without a known cell-type specific topography, employing SCAMPR successfully identified unique cell-type specific changes in gene expression due to ELS. Furthermore, we determined that the type of response to ELS varied across cell subtypes, with some of the changes due to less mean expression of a gene across a cell-type (*Gpr65*), while others were reflected in a decrease in the number of neurons comprising a cell-type that expressed a second target gene (*Pvalb*) (Figure S2, 6D, 6F). In summary, SCAMPR also provides a cost-effective approach to high resolution detection of potential, single-cell differences in gene expression in experimental models, with spatial details retained.

### Limitations of Study

SCAMPR, which utilizes HuC/D signal for cellular segmentation and ROI generation, is currently limited to the study of neurons in the nervous system. While the stability of reagents under multiplex *in situ* hybridization conditions would need to be assessed, it is likely that antibodies detecting proteins to mark glial cells or cells involved for neural vasculature and immune function can be utilized with SCAMPR to assay gene expression in non-neuronal cell populations. In addition, cell-type-specific antibodies can be validated in non-neural normal or pathogenic tissues, organoids and other sources of tissue sections and used with SCAMPR to assay gene expression spatially.

The SCAMPR pipeline utilizes three cellular attributes — gene expression, soma size, and cellular topography — to distinguish neuron populations. Yet, neuronal innervation patterns, often in combination with the three attributes utilized by SCAMPR, are also important for categorizing neurons into distinct subtypes. It has been demonstrated that one-round of RNAscope is compatible with retrograde tracers, making it feasible to incorporate a fourth modality, circuit topology, into the SCAMPR pipeline (Rana et al., 2020). For this to be successfully incorporated with multi-round HiPlex RNAscope, the signal from the traced cells would require bleaching or degradation after an initial round of imaging, making the microscopy channel available for use in the proceeding rounds to assay mRNA expression.

## Acknowledgments

We are grateful to Dr. Esteban Fernandez and The Saban Research Institute (TSRI) Cell Imaging Core for advice on image analysis methods and microscopy support, and Dr. Rand Wilcox, Dr. Ramon Durazo-Arvizu, and The Saban Research Institute Biostatistics Core for biostatistical consultation. The research was supported by a TSRI Pre-Doctoral Fellowship (RAG), NIMH grant MH067842, NIDDK grant DK126085, and the Simms/Mann Family Foundation Chair in Developmental Neurogenetics (PL).

## Author Contributions

RG and VM conceived of and developed the data analysis methodology, RG, VM, AK and PL provided recommendations for modification of the studies, AK generated the experimental material using the ELS model, RG wrote the paper and VM, AK and PL edited the manuscript.

## STAR Methods

### KEY RESOURCES TABLE

| REAGENT or RESOURCE | SOURCE | IDENTIFIER |
| --- | --- | --- |
| **Antibodies** |  |  |
| HuC/HuD | ThermoFisher Scientific | A-21271 |
| Alexa Fluor 594 AffiniPure F(ab')<sub>2</sub> Fragment Donkey Anti-Mouse IgG (H+L) | Jackson ImmunoResearch | 715-586-151 |
| **Chemicals, peptides, and recombinant proteins** |  |  |
| Tissue-Tek Optimal Cutting Temperature Compound | Sakura Finetek | 4583 |
| ProLong Gold Antifade Mountant | Jackson ImmunoResearch | Cat. #017-000-121 |
| Tween-20 | Sigma-Aldrich | P2287 |
| SSC | G. Biosciences | 786-023 |
| Bovine Serum Albumin | Gemini Bio | 700-106P |
| Normal Donkey Serum | Jackson ImmunoResearch | 017-000-121 |
| **Critical commercial assays** |  |  |
| RNAscope Intro Pack for HiPlex12 Reagent Kit – Fresh/Fixed Frozen – Mm | ACD Biosciences | 324102 |
| RNAscope HiPlex Probe – Mm -- Pvalb – T1 | ACD Biosciences | 421931-T1 |
| RNAscope HiPlex Probe – Mm -- Met – T2 | ACD Biosciences | 405301-T2 |
| RNAscope HiPlex Probe – Mm – Piezo1 – T3 | ACD Biosciences | 400181-T3 |
| RNAscope HiPlex Probe – Mm -- Chat– T4 | ACD Biosciences | 408731-T4 |
| RNAscope HiPlex Probe – Mm – Foxo1 – T5 | ACD Biosciences | 485761-T5 |
| RNAscope HiPlex Probe – Mm – Slc17a7 – T6 | ACD Biosciences | 416631-T6 |
| RNAscope HiPlex Probe – Mm – Cdh13 – T7 | ACD Biosciences | 443251-T7 |
| RNAscope HiPlex Probe – Mm – Qrfpr – T8 | ACD Biosciences | 566341-T8 |
| RNAscope HiPlex Probe – Mm – Mab21l1 – T9 | ACD Biosciences | 485051-T9 |
| RNAscope HiPlex Probe – Mm – Vip – T10 | ACD Biosciences | 415961-T10 |
| RNAscope HiPlex Probe – Mm – Gfra1 – T11 | ACD Biosciences | 431781-T11 |
| RNAscope HiPlex Probe – Mm – Cckar – T12 | ACD Biosciences | 313751-T12 |
| RNAscope HiPlex Probe – Mm -- Ntsr1 – T2 | ACD Biosciences | 422411-T2 |
| RNAscope HiPlex Probe – Mm -- Piezo1 – T3 | ACD Biosciences | 400181-T3 |
| RNAscope HiPlex Probe – Mm -- Npy2r – T4 | ACD Biosciences | 315951-T4 |
| RNAscope HiPlex Probe – Mm -- Htr3b – T5 | ACD Biosciences | 497541-T5 |
| RNAscope HiPlex Probe – Mm -- Trpv1 – T6 | ACD Biosciences | 313331-T6 |
| RNAscope HiPlex Probe – Mm -- Scn1a – T7 | ACD Biosciences | 434181-T7 |
| RNAscope HiPlex Probe – Mm -- Scn10a – T8 | ACD Biosciences | 426011-T8 |
| RNAscope HiPlex Probe – Mm -- Glp1r – T9 | ACD Biosciences | 418851-T9 |
| RNAscope HiPlex Probe – Mm -- Gpr65 – T10 | ACD Biosciences | 431431-T10 |
| RNAscope HiPlex Probe – Mm -- Piezo2 – T11 | ACD Biosciences | 400191-T11 |
| **Experimental models: Organisms/strains** |  |  |
| C57BL/6J Mice | The Jackson Laboratory | 000664 |
| **Software and algorithms** |  |  |
| SCAMPR_FilenamesCSV | GitHub | https://zenodo.org/badge/latestdoi/443220284 |
| SCAMPR_AreaFraction ImageJ Macro | GitHub | https://zenodo.org/badge/latestdoi/443220284 |
| SCAMPR_ViolinPlots | GitHub | https://zenodo.org/badge/latestdoi/443220284 |
| SCAMPR_ExpressionTopomaps | GitHub | https://zenodo.org/badge/latestdoi/443220284 |
| SCAMPR_LoessCoexpressionTopomaps | GitHub | https://zenodo.org/badge/latestdoi/443220284 |
| SCAMPR_CorrelationCirclePlots | GitHub | https://zenodo.org/badge/latestdoi/443220284 |
| SCAMPR_hClustHeatmapTopomaps | GitHub | https://zenodo.org/badge/latestdoi/443220284 |
| SCAMPR_hClustViolinPlotsCellAreas | GitHub | https://zenodo.org/badge/latestdoi/443220284 |
| SCAMPR_LoessScatterplotComparisons | GitHub | https://zenodo.org/badge/latestdoi/443220284 |
| SCAMPR_MeanComparisonsCellTypes | GitHub | https://zenodo.org/badge/latestdoi/443220284 |
| SCAMPR_PercentCellsComparisonsCellTypes | GitHub | https://zenodo.org/badge/latestdoi/443220284 |
| SCAMPR_CorrelationRegressionComparisonsMatrix | GitHub | https://zenodo.org/badge/latestdoi/443220284 |
| SCAMPR_MeanComparisonsMatrix | GitHub | https://zenodo.org/badge/latestdoi/443220284 |
| SCAMPR_PercentCellsComparisonsMatrix | GitHub | https://zenodo.org/badge/latestdoi/443220284 |

#### Lead contact

Further information help with code should be directed to the lead contact, Ramin Ali Marandi Ghoddousi.

#### Materials availability

This study did not generate new unique reagents or mouse lines.

#### Data and code availability

HiPlex Gene-Count matrices and all code used to analyze each Gene-Count matrix are publicly available at GitHub as of the date of publication (https://github.com/ramin-ali-marandi-ghoddousi/SCAMPR). DOI numbers are listed in the key resources table. Microscopy data, HiPlex images, and any additional information required to reanalyze the data reported in this paper will be shared by the lead contact upon request.

#### Experimental models and subject details

##### Animals

One female C57BL/6J mouse, sacrificed at postnatal day 100 (P100), was used for the V1 cortex HiPlex RNAscope experiments. Four ELS and four CAU litters were used for the ELS experiments. From each litter, 1 male and 1 female postnatal day 9 (P9) C57BL/6J mouse was chosen at random and sacrificed for the NG HiPlex RNAscope experiments.

All experimental procedures were performed in accordance with the Institutional Animal Care and Use Committee of The Saban Research Institute, Children’s Hospital Los Angeles.

Any additional information required to reanalyze the data reported in this paper is available from the lead contact upon request.

### METHOD DETAILS

#### Sequential Steps for SCAMPR Pipeline

##### Tissue collection and preparation

Prior to collection of adult visual cortex and early postnatal nodose ganglion, mice were anesthetized with a Ketamine:Xylazine mixture (100 mg/kg: 10 mg/kg) and transcardially perfused with 4% paraformaldehyde (pH 7.4). Due to the small size of the tissue, a dissecting microscope was used to collect the nodose ganglion from each mouse. All collected tissue was postfixed in 4% paraformaldehyde for 2-3 hours at 4°C, then cryoprotected overnight in 20% sucrose. Tissue samples were embedded in Tissue-Tek Optimal Cutting Temperature Compound (OCT) (Sakura Finetek, Cat. #4583), frozen over dry ice, and stored at -80°C. 20 µM cryosections were collected in six series through the entirety of the nodose ganglion and mounted directly onto slides. For the visual cortex, 20 µM cryosections were collected in the sagittal orientation in 4 series. All slides were stored at -20°C until processing.

##### Dual Immunohistochemistry-HiPlex RNAscope and Imaging

The RNAscope HiPlex Assay was performed according to the manufacturer’s standard protocol using the RNAscope HiPlex12 Kit (Advanced Cell Diagnostics Cat. #324194). Tissue sections were baked for 1 hour at 60°C and dehydrated in an ethanol series, followed by antigen retrieval (5 minutes at 100°C) and protease treatment (Protease IV for 30 minutes at room temperature for NG, Protease III for 30 minutes at 40° C for V1). Probes for twelve target genes were hybridized for 2 hours at 40°C, washed, and hybridized with target-binding amplifiers allowing for signal amplification of single RNA transcripts. Hybridization with negative control probes targeting bacterial genes was performed in parallel. The final step of the first round of hybridization attached three fluorophores to the first three of twelve of the target genes (T1-T3). Once the fluorophores were hybridized to the three genes, the sections were counterstained with DAPI for 30 seconds, then mounted/coverslipped with ProLong Gold Antifade Mountant (Jackson ImmunoResearch, Cat. #017-000-121). For the NG samples, signal detection was performed in four rounds, where three target genes were labeled with three cleavable fluorophores and imaged with a 40X water objective each round on a Zeiss LSM 710 confocal microscope. Experimenters were blind to the genotype of all animals during imaging and signal quantification of the NG sections. For the V1 sample, signal detection was performed in three rounds (4 probes each round) due to the availability of an infrared fluorophore (ACD Biosciences Catalog No. 322830) and a Leica Stellaris 5 confocal microscope with infrared detector. The V1 sections were imaged using a 40x water objective on a Leica Stellaris 5 and automated, inter-tile registration and blending was performed for the visual cortex images using the Stellaris imaging software (LAS X 4.1) to resolve any coordinate discrepancies between adjoining tiles.

For each section, the gain and laser power were qualitatively optimized by the experimenter for each channel (DAPI, Alexa Fluor 488, Atto 550, Atto 647N, AF750 – V1 only) and a 20 µM z-stack image was obtained (1.33 µM step-size). No signal was observed in the negative control tissue sections, or after cleaving all fluorophores, demonstrating the specificity of the assay.

After the sections were imaged, the coverslips were removed in 4X SSC buffer (G. Biosciences, Cat. #786-023) and the fluorophores were cleaved using the cleaving solution provided in the kit. A new set of fluorophores targeting the next three genes (T4-T6 for NG, T5-T9 for V1) were hybridized onto the tissue sections, another round of DAPI counterstaining was performed, and the sections were reimaged as described above. This was repeated until all 12 target genes were imaged. In order to identify the neurons in the NG and V1, immunofluorescence labeling was performed in the same tissue sections following the cleaving of the fluorophores from the last round of the HiPlex Assay.

The coverslips were washed off once again, tissue sections were briefly washed in 0.005% Tween-20 (Sigma-Aldrich Cat. #P2287) in PBS before incubation for 30 minutes in blocking solution containing 10% normal donkey serum (Jackson ImmunoResearch Cat. #017-000-121) and 1% bovine serum albumin (Gemini Bio Cat. #700-106P) in PBS. Slides were incubated overnight at room temperature with antibodies against the neuronal marker HuC/HuD (1:500, ThermoFisher Scientific Cat. #A-21271, RRID:AB_221448) with 1% BSA in PBS. Slides were washed in 0.005% PBST, then incubated for 1 hour at room temperature in Alexa Fluor 594 AffiniPure F(ab’)_2_ Fragment Donkey Anti-Mouse IgG (H+L) (1:500, Jackson ImmunoResearch Cat. #715-586-151, RRID:AB_2340858) with 1% BSA in PBS. Following three washes in 0.005% PBST and one wash in PBS, slides were counterstained with DAPI (Advanced Cell Diagnostics Cat. #324108) and coverslipped using ProLong Gold Antifade Mountant (ThermoFisher Scientific Cat. #36930). One final round of imaging was performed as described above to capture the HuC/D and DAPI signals.

##### Image preprocessing

Each confocal LSM (NG) or LIF (V1) image file was loaded into FIJI ImageJ. A maximum intensity z-projection was created, each channel (corresponding to a specific gene / HuC/D / DAPI) was split into a separate image file, and each new image was saved as a TIFF file. Generating flattened z-projections at this step provides **major advantages to efficiency** in the downstream processing; image registration and cell segmentation only needs to be performed once for each z-projected image, in contrast to separate registration and segmentation of each of the multiple z-planes in each image. To increase the speed of this step, a macro was generated in FIJI ImageJ to create and save maximum intensity z-projections for all images in a folder or set of subfolders.

Next, images from multiple rounds of imaging the same tissue were registered together using the ACD Biosciences Image Registration Software, following the manufacturers protocol (ACD Biosciences Document No. 30065UM). In brief, a DAPI image from one of the rounds of imaging was used as the reference image. DAPI images from all other rounds of imaging were registered to this reference image, generating a transformation matrix of coordinate conversions that was applied to the remaining gene and HuC/D images from each imaging round to create one, unified coordinate system for all images from all rounds. In the same software, non-overlapping regions around the edges of the images are cropped, and each registered image is saved separately as a TIFF file for downstream processing. An alternative registration method using ImageJ is detailed on the SCAMPR GitHub.

#### Cell Segmentation and Signal Quantification

Manual cell segmentation was performed by opening all registered HuC/D images in FIJI ImageJ, then using the freehand selection tool to circle individual cells, which were each outlined as separate regions of interest (ROIs). ROIs for each image were then saved into a ZIP file for use in later processing. Automated cell segmentation was performed using Cellpose (Stringer et al., 2021). A Google Colab notebook that was provided by the authors of Cellpose was modified and used to generate the ROIs for each image based on the HuC/D images. The ROI outlines that are generated by Cellpose were converted into FIJI ImageJ compatible ROIs using the Cellpose_Outline_to_ROI_Converter.py script and saved as a ZIP file for use in later processing.

Semi-automated signal quantification using SCAMPR requires limited user inputs. In ImageJ, background subtraction was used to remove imaging artifacts such as autofluorescence and other noise. Next, global image thresholding was performed to designate each pixel in the image as either signal or background. Both steps require the selection of specific parameters: a rolling-ball radius pixel size for the background subtraction, and a minimum threshold value for the global image thresholding. Due differential tissue quality, imaging, and mRNA expression levels between multiple genes, one set of parameters does a poor job of separating signal from background for all mRNAs across multiple tissues. For this reason, experimenter-blinded manual selection of the rolling-ball radius and minimum threshold value is required for each image prior to signal quantification. The rolling-ball radiuses and threshold values for image files of interest are saved in one CSV file, which is utilized by the FIJI ImageJ macros that perform automated signal quantification in downstream steps.

A FIJI ImageJ macro was developed as part of SCAMPR to quantify whole cell mRNA signal from RNAscope experiments. The inputs for this macro, which were generated and processed in previous steps, are as follows: 1) the maximum intensity z-projected images each gene, 2) the cellular ROIs generated manually or through utilization of Cellpose, and 3) the CSV file containing the optimal, image-specific rolling-ball radius and minimum threshold as determined by the experimenter. Using this information, these macros determine the expression of all genes in all cells for the input images, thereby generating a Gene by Cell matrix file with the results. Each row of this matrix corresponds to a single cell and contains the animal ID, the image ID, the ROI (cell) area, and the expression level of each gene of interest. The gene expression level is displayed as an area-fraction calculation (the number of pixels expressing the gene within the cell divided by the area of the cell in pixels). This area-fraction expression value can be used as-is for the downstream analysis or can be converted to a value representing the number of pixels expressing the gene by multiplying the area-fraction by the cell area (i.e. pixel coverage in a cell).

#### Quantitative Analysis and Spatial Mapping

The first step prior to analysis is to import the Gene by Cell matrix created in the previous step into RStudio. Cells not expressing any genes can be removed from the dataset if desired. Depending on the genes chosen for the experiment, this can help remove noise due to misclassification of tissue imperfections or other structures as cells. Next, the columns containing the gene-expression data can be left as area-fraction calculations or converted into pixel coverage. These values are then natural log normalized to reduce high-expressing gene bias during analysis and data visualization and used to generate violin plots, co-expression charts, spatial maps, and cluster heatmaps/dendrograms.

##### Violin plots

Violin plots can be generated in two ways: 1) by plotting the gene expression levels of each gene in each cell, or 2) by only plotting the expression levels of genes after excluding cells that do not express the gene of interest. The latter provides a comparison of gene expression levels, while the former will give a comparison of both expression levels as well as a relative comparison of the number of cells that express a particular gene.

##### Expression, co-expression, and spatial mapping

Cellular ROIs are uploaded into R and matched to their appropriate cells in the Gene by Cell matrix. If desired, manual annotations that mark structural boundaries (e.g the layers in the V1) or other image features are also generated in ImageJ and uploaded into R. The gene expression levels for a gene of interest are binned into different expression ranges and added as a column to the matrix. Each bin is assigned a color, and these assigned colors are also appended to the matrix. Each cell is then plotted based on its ROI coordinates and colored according to its gene expression bin.

Graphical representation of gene co-expression is generated in R by using the log normalized gene expression values. Correlation coefficient matrices can be plotted alone or combined with ordinary least squares regression lines for each gene pair. For these matrices, the data can be stratified based on group, making it useful for exploratory analysis of differences in gene co-expression between groups.

For co-expression spatial mapping, a local polynomial regression model (LOESS) is fitted to the log normalized gene expression values of two genes of interest. Residual distances are calculated for each cell then binned based on size, and a color code from a red-blue scale is assigned to each bin. This color code is added as a column to the Gene by Cell matrix. Cells are plotted according to their ROI coordinates, and the new matrix is then used to color each cell according to its residual size bin.

##### Clustering and heatmaps

Log normalized data are hierarchically clustered. Depending on the dataset, different distance measures (Euclidean, Pearson, and Spearman) can be utilized as inputs into the clustering algorithm, and a choice can be made between multiple agglomeration methods (Ward.d2, complete, etc.). Here, we employed Ward.d2 clustering on Spearman correlation coefficient distance measures. Cluster identities are assigned to each cell, and heatmaps are generated using normalized gene expression data. The rows (cells) in the heatmap are organized by hierarchical clusters.

Cellular cluster identities are appended to the Gene by Cell matrix and used in conjunction with ROIs to spatially map cells based on cluster. Cluster identities also can be utilized in conjunction with gene expression levels and cell area to generate violin and boxplots, allowing one to look for gene expression differences between clusters.

#### Early Life Stress Paradigm

A limited bedding and nesting (LBN) model was used to expose pups in our experimental group to chronic early life stress (ELS) from P2 to P9 (Peña et al., 2017). In brief, each ELS cage is lined with a mesh platform and just enough bedding is placed under the platform to sparsely cover the cage floor. The dam is provided with half of standard nesting material. Combined limited bedding and nesting materials have been shown to act as a stressor on the dam, and consequently leads to fragmented and erratic maternal care of the pups. Care-as-usual (CAU), control litters are raised in cages with normal bedding and nesting resources. At P9, one male and one female mouse were chosen at random from each of 4 different litters and sacrificed for HiPlex experiments.

12 genes that are involved in classifying NG neurons into 18 putative functionally projecting cell-types were compared between groups in these experiments (Kupari et al., 2019).

#### Manual and Cellpose Segmentation Comparisons

Eight 384 x 344 pixel ROIs, containing a combined total of 307 cells with a mean of 38 cells per ROI, were randomly cropped from different locations in the visual cortex image and were used to assess the time demand and accuracy associated with manual segmentation, automated segmentation using Cellpose, and semi-automated segmentation by combining Cellpose and manual correction.

##### Time requirement experiments

To determine the time required for manual segmentation, each image was opened in ImageJ, and the amount of time needed to segment each cell (ROI) by hand was determined. For these experiments, each cell was segmented using a stylus on a touchscreen laptop or tablet in combination with the free hand selection tool on ImageJ. For Cellpose segmentation, the amount of time required to load all eight files into the Cellpose Google Colab notebook, to run the program, and to output a ZIP file of all ROI masks was calculated. For the Corrected Cellpose time calculation, Cellpose ROIs were overlayed onto each image in ImageJ, and the amount of time needed to delete inaccurate Cellpose ROIs and redraw boundaries by hand using the free hand selection tool was calculated. To determine the total time required for Corrected Cellpose segmentation, the time to perform ROI corrections was added to the average time for segmentation using Cellpose alone.

##### Accuracy Experiments

Segmentation accuracy was calculated by comparing the ROIs from Cellpose and Corrected Cellpose to the ROIs from the manual segmentation. Using the centroid of each ROI, the nearest Cellpose or Cellpose Corrected neighbor for each Manual ROI was determined in ImageJ. The percentage of overlap was calculated for each neighboring pair. In R, the number of true positive (TPs), false positives (FPs), and false negatives (FNs) was determined at a 70% overlap threshold. The F1 score was calculated by using the following equation:

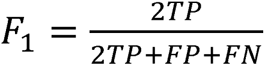

### QUANTIFICATION AND STATISTICAL ANALYSIS

All statistics were performed in R and are described in the results section as well as the figure legends. In the segmentation method comparisons, n represents the number of images that were used for the time and accuracy calculations in each group. In the ELS NG comparison experiments, n represents the number of animals per group. For all comparisons, means were compared using a Student’s T test and significance was reached at p < 0.05.

## Figure Legends

**Supplemental Figure 1:**
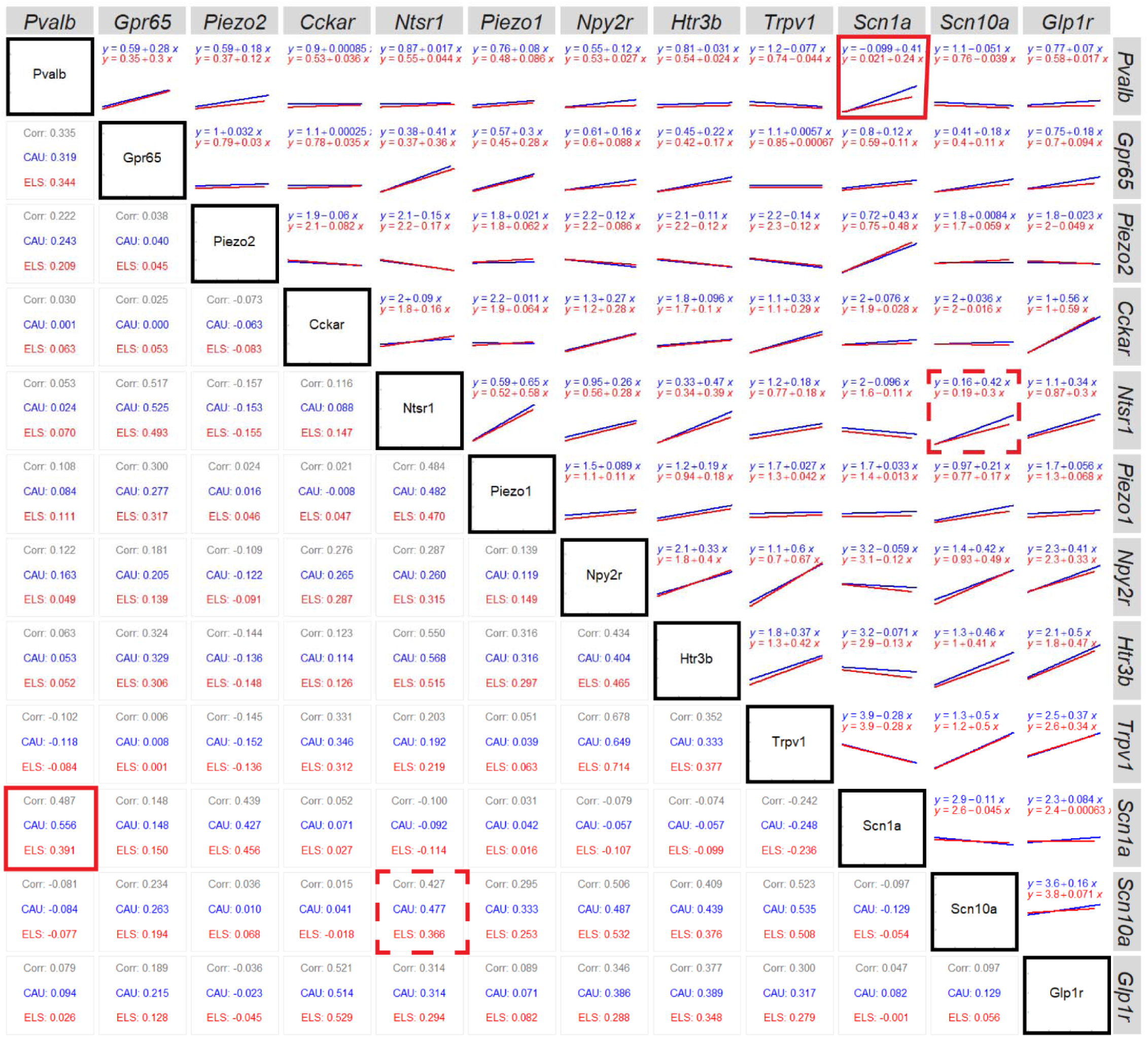
Pairwise gene co-expression comparisons between CAU and ELS in nodose ganglion. In the bottom-left of the diagonal, Pearson’s correlation coefficients were calculated for the expression of each gene pair for the mice in the CAU group (blue), for the mice in the ELS group (red), and for all mice from both groups (grey). In the top-right of the diagonal, ordinary least squares regression models were fitted to the expression of each gene pair in the CAU (blue) and ELS (red) mice. The equation for each regression line is also plotted. Red boxes indicate gene pairs that exhibited sizable differences in correlation coefficients and regression slopes between groups and were selected for further analysis using modern statistical methods.

**Supplemental Figure 2:**
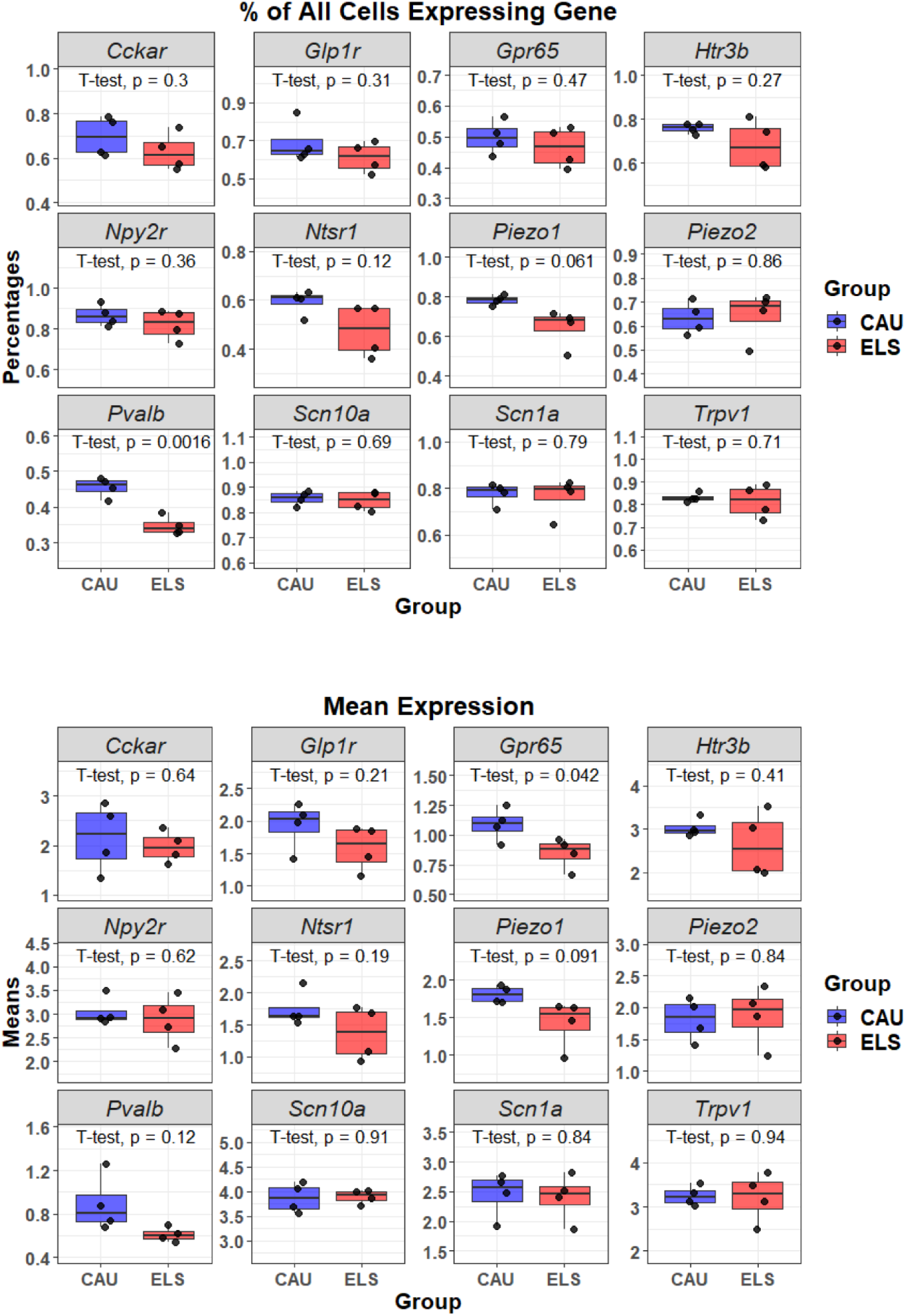
Comparison of gene expression between CAU and ELS nodose ganglia neurons. **A.** The percent of cells expressing each gene in each animal in the CAU group are averaged and compared to the averages calculated for the ELS animals. **B.** Mean gene expression in each animal in the CAU group is calculated and compared to the mean gene expression in the ELS group. Student’s t-tests are performed to compare means between groups and p-values are calculated to assess significance. Note the significant decrease in the percentage of all cells expressing *Pvalb*, but no change in mean expression level. Conversely, the percentage of *Gpr65* neurons is unchanged between CAU and ELS groups, but the mean expression level is decreased in the ELS group.

